# A FISH-based chromosome map for the European corn borers yields insights into ancient chromosomal fusions in the silkworm

**DOI:** 10.1101/013284

**Authors:** Yuji Yasukochi, Mizuki Ohno, Fukashi Shibata, Akiya Jouraku, Ryo Nakano, Yukio Ishikawa, Ken Sahara

**Author notes:** Note: Nucleotide sequence data reported are available in the DDBJ/EMBL/GenBank databases under the accession numbers LC002981–LC003010. Corresponding author: Yuji Yasukochi, Owashi 1-2, Tsukuba, Ibaraki 305-8634, Japan.

## Abstract

A significant feature of the genomes of Lepidoptera, butterflies and moths, is the high conservation of chromosome organization. Recent remarkable progress in genome sequencing of Lepidoptera has revealed that syntenic gene order is extensively conserved across phylogenetically distant species. The ancestral karyotype of Lepidoptera is thought to be *n* = 31; however, that of the most well studied moth, *Bombyx mori*, is *n* = 28, suggesting that three chromosomal fusion events occurred in this lineage. To identify the boundaries between predicted ancient fusions involving *B. mori* chromosomes 11, 23 and 24, we constructed FISELbased chromosome maps of the European corn borer, *Ostrinia nubilalis* (*n* = 31). We first determined 511 Mb genomic sequence of the Asian corn borer, *Ostrinia furnacalis*, a congener of *O. nubilalis*, and isolated BAC and fosmid clones that were expected to localize in candidate regions for the boundaries using these sequences. Combined with FISH and genetic analysis, we narrowed down the candidate regions to 40kb – 1.5Mb, in strong agreement with a previous estimate based on the genome of a butterfly, *Melitaea cinxia*. The significant difference in the lengths of the candidate regions where no functional genes were observed may reflect the evolutionary time after fusion events.

## Introduction

Accumulating evidence currently reveals that the gene order and content of each chromosome are highly conserved among lepidopteran species, butterflies and moths. Since we first described syntenic gene order on four pairs of chromosomes between the silkworm, *Bombyx mori* and a butterfly, *Heliconius melpomene* (Yasukochi *et al.* 2006), a number of reports have demonstrated extensive synteny between *B. mori* and other species based on genetic linkage studies (Pringle *et al.* 2007; Beldade *et al.* 2009; Baxter *et al.* 2011; Van't Hof *et al.* 2012) and chromosomal FISH (fluorescence *in situ* hybridization) analysis (Yasukochi *et al* 2009; Yoshido *et al.* 2011; Sahara *et al.* 2013).

Genome sequencing with high resolution assignments to chromosomes enabled more direct and precise comparison between species (International Silkworm Genome Consortium 2008; *Heliconius* Genome Consortium 2012; Ahola *et al.* 2014), and revealed that the genomes of *B. mori* (*n* = 28) and *H. melpomene* (*n* = 21) are composed of 31 well-conserved chromosomal segments corresponding to 31 chromosomes of another butterfly, *Melitaea cinxia* (Ahola *et al.* 2014). Since *n* = 31 is the most common chromosome number observed in distantly related families of Lepidoptera (Robinson 1971), it is likely that *n* = 31 was the karyotype of the common ancestor and chromosomal fusion events occurred in the lineages leading to *B. mori* and *H. melpomene* (*Heliconius* Genome Consortium 2012; Ahola *et al.* 2014). This hypothesis would be greatly strengthened if other species of *n* = 31 karyotype which are distantly related to *M. cinxia* are shown to have similarly organized chromosomes.

The genus, *Ostrinia*, belonging to the superfamily Pyraloidea, which is closely related to neither *B. mori* nor butterflies (Regier et al. 2013), is widely utilized for analyzing evolution in female sex pheromone biosynthesis and male recognition systems during spéciation (for review, see Lassance 2010). Among them, the Asian and European corn borers, *Ostrinia furnacalis* (*n* = 31) and *O. nubilalis* (*n* = 31), are serious pests of maize. Although a linkage map was constructed for *O. nubilalis* based on AFLPs and microsatellites, not many genes were mapped on it (Dopman et al. 2004). Fine mapping of *O. nubilalis* genes is useful not only for characterization of the genetical basis of complex phenotypes but also for verification of synteny. We previously succeeded in identifying three chromosomes of *O. nubilalis* orthologous to *B. mori* chromosomes 1(Z), 16 and 23 by FISH analysis (Yasukochi *et al.* 2011a).

Here, we describe construction of a cytogenetic map covering all 31 chromosomes of *O. nubilalis* using 123 BACs (bacterial artificial chromosomes) and eight fosmid clones as FISH probes. We then focused this study on the identification of boundaries between ancient chromosomes within *B. mori* chromosomes 11, 23, and 24.

## Materials and Methods

### Insects

*O. nubilalis* used in this study was previously described (Yasukochi *et al.* 2011). *O. furnacalis*, *O. scapulalis*, *O. latipennis*, *O. palustralis* and *Pleuroptya ruralis* were from wild populations collected in the Kanto area, Japan. These were fed an artificial diet, Insecta LFS (Nosan, Yokohama, Japan), in plastic cups. *B. mori* strain p50 was used and reared on mulberry leaves.

### Genomic sequencing of *Ostrinia furnacalis*

Genomic DNA was prepared from an *O. furnacalis* male pupa that was a subcultured offspring of wild insects originally collected in Inawashiro, Fukushima, Japan, using a DNeasy Blood & Tissue Kit (Qiagen, Venlo, Netherland) according to the manufacturer’s protocol. The genomic DNA was then submitted to Otogenetics Corporation (Norcross, GA, USA) for sequencing. An Illumina paired-end library with an average insert size of 196 bp was made from fragmented DNA using NEBNext reagents (New England Biolabs, Ipswich, MA, USA), and sequenced on an Illumina HiSeq 2000 which generated paired-end reads of 100 nucleotides.

### Genome Assembly

The raw paired-end reads were trimmed and filtered by Trimmomatic (Bolger et al. 2014) and reads of low quality and those contaminated with adapter sequences (approximately 15%) were removed. Then, genome assembly and scaffolding were carried out using SOAPdenovo2 (version r240) (Luo *et al.* 2012) with k-mer length 75.

### Construction of an *Ostrinia iurnacalis* fosmid library

A fosmid library was constructed by Takara Bio (Kyoto, Japan). Genomic DNA was purified from whole male pupae of *O. furnacalis* by the CTAB method (Murray and Thompson, 1980). The DNA was fragmented by sonication and fractionated by pulse-field electrophoresis. Genomic fragments ranging from 33–48kb were purified from agarose gels and cloned into a pCC1FOS vector (Epicentre, Madison, WI). A total of 75,264 single colonies were arrayed into 196 384-well microplates.

### Screening of BAC and fosmid libraries

An *O. nubilalis* BAC library, ON_Ba, was obtained from the Clemson University Genomics Institute (Clemson, SC, USA). Construction of a *B. mori* BAC library is described elsewhere (Wu et al. 1999). Detailed methods for PCR screening of large-insert libraries and FISH analysis are described in Yoshido *et al.* (2014). Briefly, the first screening was carried out against DNA pools representing each microplate containing 384 clones. A second screening was then carried out against DNA pools prepared from the mixture of cell cultures located in the same column or row of each positive microplate. This was followed by single colony isolation from frozen stocks of candidate clones and overnight cultures of the isolated clones for final PCR confirmation. PCR conditions were as follows: 3-min denaturation at 94°, followed by 45 cycles with a 1-min denaturation at 94°, 2-min annealing at 55°, and 3-min elongation at 72°, ending with a 5-min final extension at 72°. The sequences of primers used in this study are listed in Table S1.

### FISH analysis

Chromosome preparation was carried out after Sahara *et al.* (1999) with slight modifications as described in Sahara *et al.* (2013). BAC- or fosmid-DNA extraction and probe labeling were done according to the descriptions in Yasukochi *et al.* (2009), Yoshido *et al.* (2011), and Sahara *et al.* (2013). A published reprobe technique for Lepidoptera (Shibata *et al.* 2009) was used to integrate results obtained by multiple hybridizations with different probes. All FISH and chromosome images were acquired with a DFC350FX CCD camera attached to a DM 6000B microscope (Leica Microsystems) and processed with Adobe Photoshop ver 7.

### Genotyping

The F_2_ progeny of *B. mori* and the BF_1_ progeny between *O. nubilalis* and *O. scapulalis* used in this study were identical to those described in previous reports (Yasukochi *et al.* 2006; Yasukochi *et al.* 2011a). Genomic DNAs were individually amplified in the same manner as library screens. The completed reactions were denatured at 95° for 5-min, annealed at 55° for 15-min to promote heteroduplex formation, and loaded on 0.67×MDE gels (Cambrex, Rockland) in 0.5×TBE to detect polymorphisms.

## Results and Discussion

### FISH analysis using previously isolated *O. nubilalis* BAC clones

We had previously isolated 163 *O. nubilalis* BAC clones containing single-copy genes which could be used as anchors for comparing genomes (Yasukochi *et al.* 2011b). Thus, we used these BACs as probes for FISH analysis of *O. nubilalis.* Subsequently, we obtained stable and reproducible signals from 113 BACs harboring 117 conserved genes between *O. nubilalis* and *B. mori* (Table S2), which could identify 29 of 31 *O. nubilalis* chromosomes (Fig. 1). The gene order on each chromosome was well conserved between *O. nubilalis* and *B. mori* with few exceptions (Fig. 1). We designated the same chromosome numbers, 1–31 excluding 11 and 30, as those of orthologous chromosomes in previously characterized species of *n* = 31 karyotype (Baxter *et al.* 2011; Van't Hof *et al.* 2012; Sahara *et al.* 2013).

**Figure 1.**
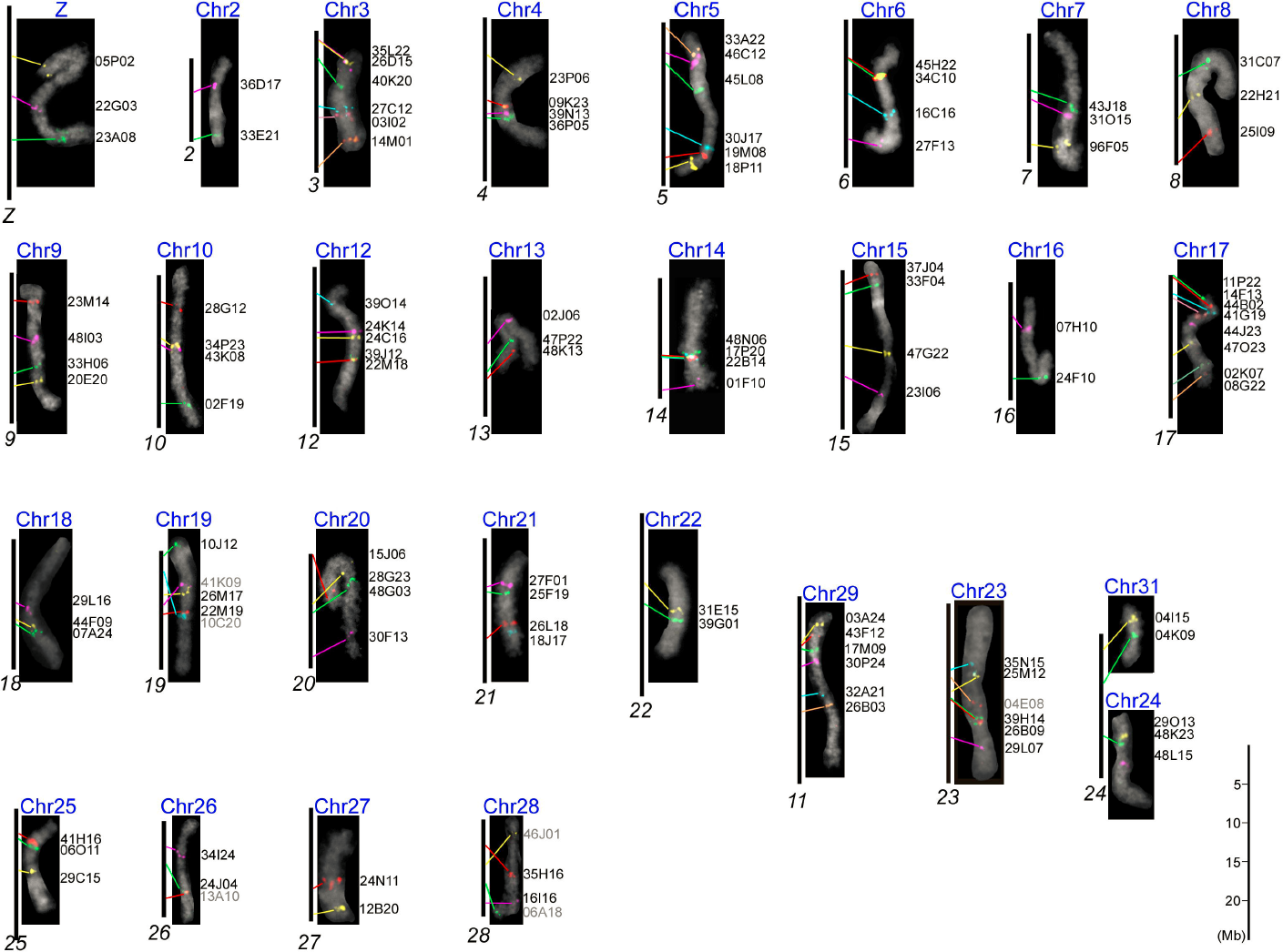
Comparison of orthologs between individual chromosomes of *Ostrinia nubilalis* and *Bombyx mori.* DAPI-stained images (gray pseudo color) of individual pachytene bivalents of *O. nubilalis* (blue chromosome numbers) show hybridization signals from *O. nubilalis* BAC probes. Black vertical bars represent orthologous *B. mori* chromosomes (italic black numbers) based on locations taken from Kaikobase (http://sgp.dna.affrc.go.jp/KAIKObase/). *O. nubilalis* BAC designations are shown on the right of each chromosome image. Gray letters indicate intrachromosomal rearrangement of localized genes and lack of conserved gene order between *O. nubilalis* and *B. mori.* Colored lines show correspondence of orthologs between *B. mori* and *O. nubilalis* based on the carrier BAC signals. Note that images of *O. nubilalis* chromosomes do not reflect true relative sizes because they were taken from different preparations.

We then investigated the possibility of performing cross-species FISH analysis since it might be useful to detect local chromosome rearrangements peculiar to certain species which might drive speciation by changes in behavior and pheromone composition. We used four *Ostrinia* species (*O. scapulalis*, *O. furnacalis*, *O. latipennis* and *O. palustralis*), as well as *Pleuroptya ruralis*, which belongs to the same subfamily, Pyraustinae. Of the four *Ostrinia* moths, *O. scapulalis* is most closely related and *O. latipennis* and *O. palustralis* are most distantly related to *O. nubilalis* (Kim *et al.* 1999). As shown in Fig. 2, using three common *O. nubilalis* BAC probes previously identified to be located on chromosome 5 we obtained clear signals in the same order on a single chromosome of these five species.

**Figure 2.**
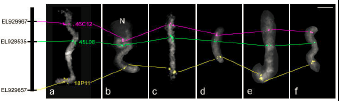
Conserved gene order on chromosome 5 of 5 *Ostrinia* species and a close relative based on cross-hybridization of 3 representative BACs from *O. nubilalis.* a. *O. nubilalis'*, b. *O. furnacalis*, c. *O. scapulalis*, d. *O. latipennis*, e. *O. palustralis*; and f. *Pleuroptya ruralis.* Signals from BAC probes 46C12, 45L08 and 18P11, which carry orthologues EL92996, EL928535 and EL929657, are pseudocolored with magenta, green and yellow, respectively. The black bar represents *B. mori*chromosome 5. Scale bar, 5μm.

### Sequencing of the *O. furnacalis* genome

As described above, we did not have any BAC probes ostensibly located on chromosomes 11 and 30. For BAC isolation, we had utilized sequences of known genes and ESTs in public databases or newly determined sequences of PCR fragments amplified by nested degenerate primers designed from conserved amino acid sequences (Yasukochi *et al* 2011b). However, because genes expressed in limited stages and tissues are rarely included in ESTs, it was very difficult to find an adequate number of conserved genes on such narrow chromosomal regions having no markers. In addition, cDNA sequences lack sequences of introns, which makes it difficult to design PCR primers and to judge whether amplified fragments are genuine or non-specific.

Thus, we decided to obtain genome sequences of an *Ostrinia* species, which would greatly facilitate finding targeted genes and establishing genetic markers in targeted locations. Since *O. nubilalis* does not occur in Japan, strict escape prevention measures are required for rearing it. Consequently, we used a local congener, *O. furnacalis*, as its substitute. We sequenced a paired-end library (average insert size 196 bp) constructed from an *O. furnacalis* male pupa and obtained 511 Mb assembled sequences comprising 745,531 scaffolds and singleton contigs (N50 = 930bp) (Table S3).

### Screening of *O. furnacalis* orthologues

Table S4 lists lepidopteran genes that were previously identified to be located near the region orthologous to the boundaries between ancient chromosomes in *B. mori* before our analysis of *O. furnacalis* genome sequences (Beldade *et al.* 2009; Yasukochi *et al.* 2009; Van't Hof *et al.* 2012; Sahara *et al.* 2013). The boundaries were predicted to be located at position 4,006,128–6,271,907 of chromosome 11, 17,071,002–17,410,092 of chromosome 23 and 8,719,098–11,527,820 of chromosome 24 in the *B. mori* genome database, Kaikobase (Shimomura *et al.* 2009). Subsequently, we obtained full-length cDNAs, gene models and known genes that were predicted to localize in these regions from Kaikobase. In addition, we selected several *B. mori* sequences located at position 1–4,006,128 of chromosome 11 and position 17,410,092–23,133,604 of chromosome 23, which correspond to unidentified *O. nubilalis* chromosomes 11 and 30.

Candidate regions of ancient chromosomal fusions were located within single scaffolds, Bm_scaf59 and 12, of *B. mori* chromosomes 11 and 23. In contrast, the candidate region in *B. mori* chromosome 24 was separated into three scaffolds. Furthermore, we could not find any conserved genes in the middle scaffold, Bm_scaf 147 (position 9,817,336–10,242,811), since no known genes or full-length cDNAs were located there and gene models and transcripts assigned to the scaffold did not show significant similarities to available *O. furnacalis* genome sequences. Ultimately, we utilized 43 *B. mori* sequences as queries for a tblatn search against the *O. furnacalis* genome sequences. Thirty of the query sequences showed striking similarities to *O. furnacalis* scaffolds and contigs containing well-conserved single-copy genes (Table l).

**Table 1.**
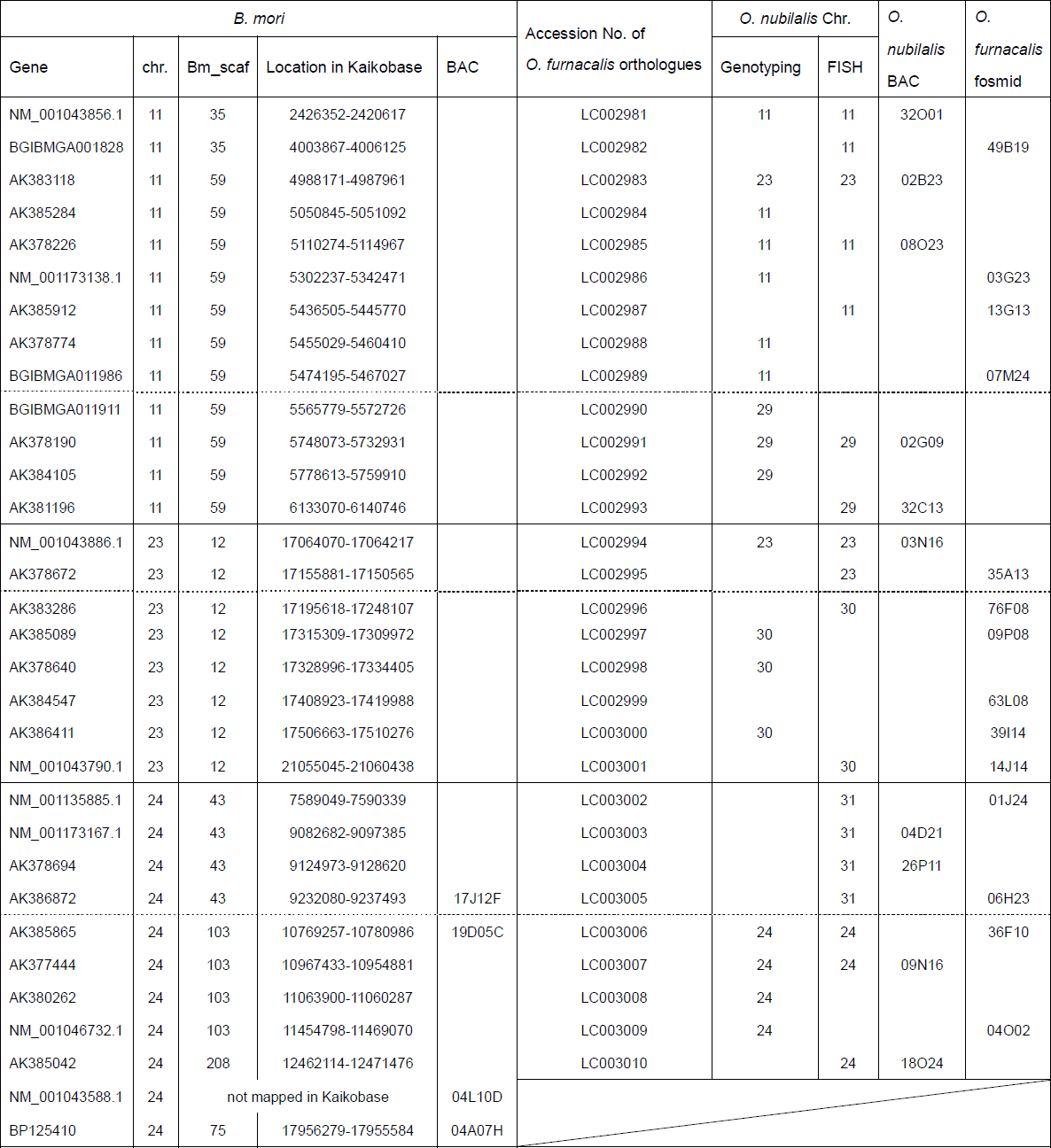
Summary of the results onbtained by FISH and genetic analysis in this study

### Screening of *O. nubilalis* BAC and *O. furnacalis* fosmid libraries

We then designed 30 pairs of PCR primers to isolate genomic clones containing candidate genes (Table S1). In general, BACs are preferable as FISH probes compared with fosmids, presumably because longer inserts can label longer chromosomal regions more stably. In addition, the chromosome preparations used for this analysis were derived from *O. nubilalis*, which might require longer regions to form stable hybrids with *O. furnacalis* probes.

Thus, we first screened the *O. nubilalis* BAC library, followed by screening of a newly constructed fosmid library from *O. furnacalis* pupae from which we obtained 75,264 colonies (average insert size, 33.1kb; total estimate size, 2,491Mb). Consequently, we isolated 10 BACs (Table 1) from the *O. nubilalis* BAC library, and then isolated 14 fosmids (Table 1) by screening the *O. furnacalis* fosmid library with markers that had failed to isolate positive *O. nubilalis* BACs.

### FISH and genetic analysis

We performed FISH analysis against *O. nubilalis* chromosomes using the 24 BAC and fosmid probes described above. In parallel with the FISH analysis, we performed genetic analysis of the candidate genes using twenty-four BF_1_ progeny between *O. nubilalis* and *O. scapulalis* to validate and complement the FISH results, since we could not isolate BACs or fosmids for six genes (Table l).

We obtained clear signals from five BACs and two fosmids containing *Ostrinia* orthologues of *B. mori* genes located on chromosome 11. Two BACs, 02G09 and 32C13, which had been assigned to previously identified *O. nubilalis* chromosome 29, and signals from two BACs (08O23, 32O01) and two fosmids (13G13, 49B19), were mapped onto the same chromosome (Fig. 3A). Thus, we designated this newly identified chromosome as *O. nubilalis* chromosome 11. In addition, we obtained polymorphic PCR products from nine orthologues that were mapped onto *O. nubilalis* chromosomes 11 or 29, as expected (Table S5, Table 1). Taking all this data together, the boundary between ancestral chromosomes 11 and 29 on *B. mori* chromosome 11 was narrowed to the region between gene models BGIBMGA011986 and BGIBMGA011911 (Table 1, Fig. 4A).

**Figure 3.**
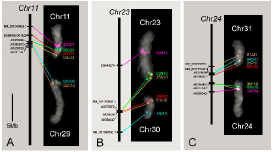
FISH detection of *O. nubilalis* chromosomes 11 and 29 (panel A), 23 and 30 (panel B) and 24 and 31 (panel C). Signals on the *O. nubilalis* chromosome (white numbers) are pseudocolored and probe names are shown to the right of the FISH images. See Table 1 for details. Left-side bars represent orthologous *B. mori chromosomes* (italic black numbers). Locations of *B. mori* orthologs are taken from Kaikobase. FISH images are from different individuals and/or preparations. Hence, *O. nubilalis* bivalent lengths vary depending on the pachytene stage.

**Figure 4.**
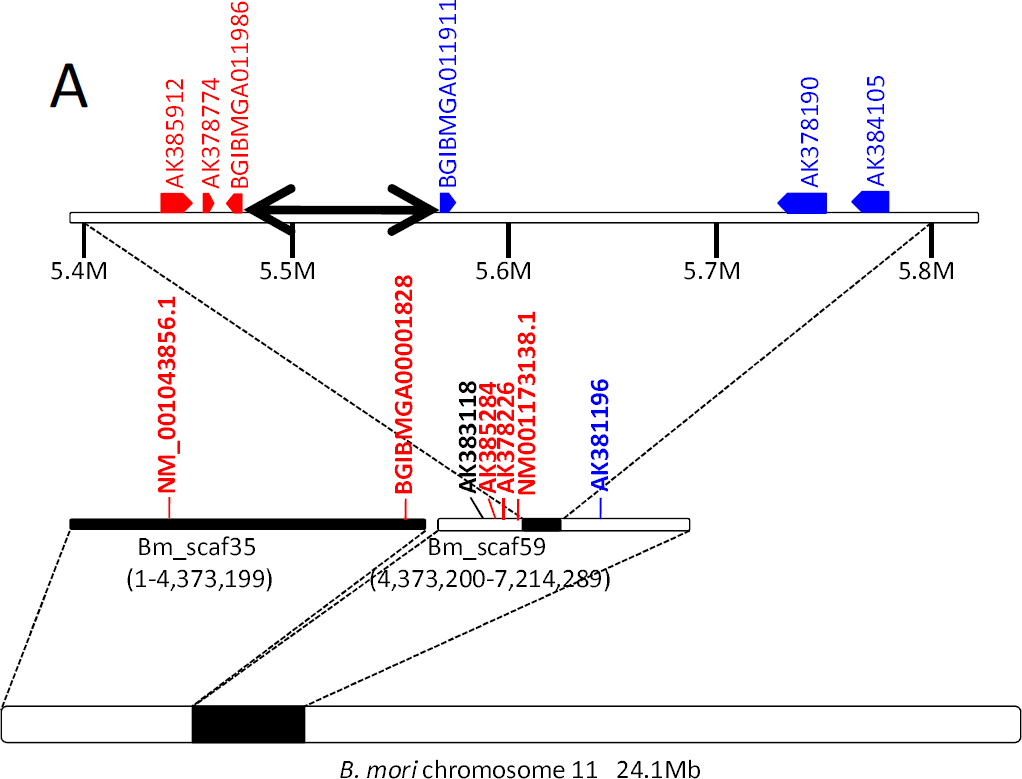
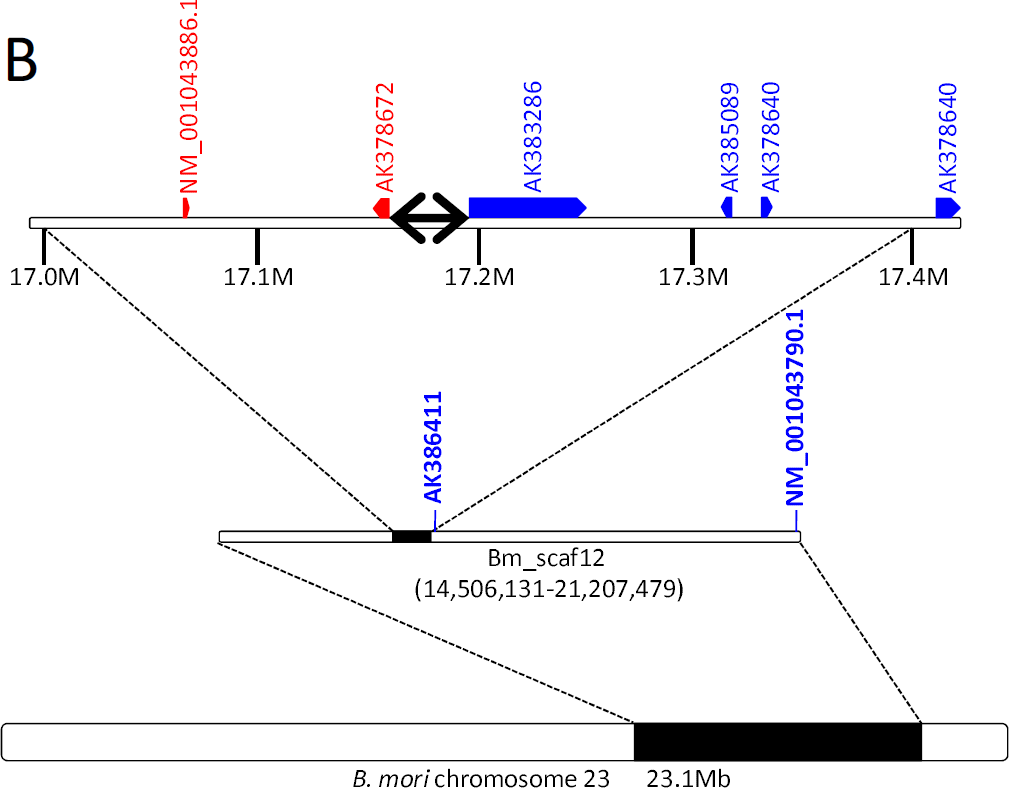
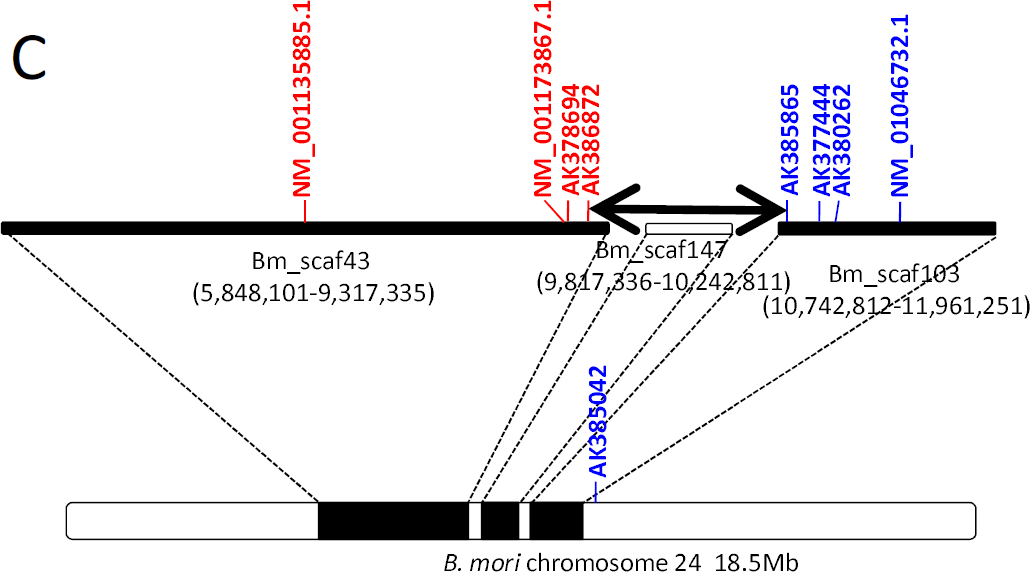
Candidate regions of ancient chromosomal fusions in *B. mori* chromosomes 11 (panel A), 23 (panel B) and 24 (panel C). Candidate regions are shown as two-headed arrows. Vertical letters indicate *B. mori* genes for which orthologues are located on *O. nubilalis* chromosomes 11 (red letters, panel A), 23 (red letters, panel B), 24 (red letters, panel C), 29 (blue letters, panel A), 30 (blue letters, panel B) and 31 (blue letters, panel C). Letters in parentheses indicate locations of *B. mori* scaffolds in Kaikobase.

Unexpectedly, BAC 02B23, which contained an *O. nubilalis* orthologue (LC002983) of a *B. mori* full-length cDNA, AK383118, was localized on another chromosome, excluding it as belonging to *O. nubilalis* chromosome 11 or 29. Genetic analysis revealed that this *O. nubilalis* orthologue was located on *O. nubilalis* chromosome 23 (Table 1, Table S5). However, there remained the possibility that AK383118 was incorrectly assembled into Bm_scaf59. Thus, we performed linkage mapping of AK383118 in *B. mori* and confirmed its location on *B. mori* chromosome 11 (data not shown). This was the only example for which we found evidence suggesting an ancient translocation.

Similarly, FISH analysis identified missing *O. nubilalis* chromosome 30 (Fig. 3B) on *B. mori* chromosome 23, where the boundary between ancestral chromosomes 23 and 30 was estimated to be localized between full-length cDNAs AK378672 and AK383286 (Table 1, Fig. 4B). Finally, *O. nubilalis* orthologues of *B. mori* genes on Bm_scaf43 and Bm_scafl03 were assigned to *O. nubilalis* chromosomes 31 and 24, respectively (Fig. 3C, Fig. 4C).

Intervals between *B. mori* scaffolds in Kaikobase are not experimentally confirmed but are designated arbitrarily as 500kb (http://sgp.dna.affrc.go.ip/KAIKObase/). Hence, we suspected that the distance between Bm_scaf43 and Bm_scafl03 was much shorter than the estimated length of 1.5Mb from the genome database. Conversely, assigning unmapped contigs located in this gap might enlarge the actual distance. Thus, we isolated *B. mori* BACs 17J12F and 19D5C containing the most distal full-length cDNAs of the scaffolds, AK386872 and AK385865, respectively (Table 1). FISH probing with these clones suggested that the interval between Bm_scaf43 and Bm_scafl03 was longer than 1Mb, although the estimate varied depending on the state of chromosome condensation (Fig. 5).

**Figure 5.**
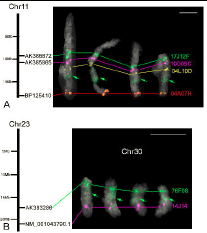
(A) A sequence gap between Bm_scaf43 and Bm_scaf 103 of *Bombyx mori* chromosome 11 revealed by BAC-FISH. Signals from 17J12F (green), 19D05C (magenta) 04L10D (yellow) and 04A07H (red) represent AK386872, AK385865, NM_001043588.1 and BP125410. Note that the green signals from 17J12F faintly stain the heterochromatic region (refer to Figure 2 in Yoshido et al 2005) of *B. mori* chromosome 11 (green arrows). Scale bars, 5μm. (B) Ectopic signals from an *O. furnacalis* fosmid (76F08) located near the end of chromosome 30 appear in *O. nubilalis* chromosome 30 (Chr30 shown in white letters). Two fosmids, 76F08 and 14J14, carrying representative single genes LC002996 and LC003001, are orthologs of *B. mori* AK383286 and NM_001043790.1, respectively (see Table 1). Black bars represent *B. mori* chromosomes 11 and 23 (Chrll and Chr23). Scale bars, 5μm.

Intriguingly, the 17J12F probe also painted a segment of chromosome 24 (Fig. 5A) which was positionally consistent with a previously described DAPI-positive and presumably heterochromatic segment (Yoshido *et al.* 2005). Similar ectopic painting was also observed for *Ostrinia* BACs and fosmids putatively located near the ends of chromosomes (Fig. 5B). A possible explanation is that these clones contain subtelomeric heterochromatin and tend to label heterochromatin scattered around chromosomes. If so, 17J12F may contain subtelomeric heterochromatin that originated from an ancient chromosomal end.

As described above, the maximum estimate of the boundary between ancient chromosomes is much longer in *B. mori* chromosome 24 (1,532kb) compared to the estimates for chromosomes 11 (91.6kb) and 23 (39.7kb) (Fig. 4). Taken together, we speculate that the fusion event generating *B. mori* chromosome 24 occurred after those generating chromosomes 11 and 23, and internal deletion of the former subtelomeric heterochromatin is still underway. Further analysis is needed to characterize karyotypes of species more closely related to *B. mori* to verify the history of chromosomal fusions in additional ancestral lineages.

### Comparison with the *Melitaea cinxia* genome data

Judging from the results described above, we estimated the positions of boundaries between ancient chromosomes in *B. mori* chromosomes 11, 23 and 24 as 5,474,195–5,565,779, 17,155,881–17,195,618 and 9,237,493–10,769,257, respectively. In the proceeding work comparing *B. mori* and *M. cinxia* genomes, these boundaries were estimated to be 5,410,160–5,798,000 in chromosome 11, 17,180,240–17,247,200 in chromosome 23 and 9,953,070–10,742,570 in chromosome 24 (Ahola *et al.* 2014). Our estimates based on FISH and genetic analysis of *O. nubilalis* conserved genes is in good accordance with those based on *M. cinxia* genome sequences in spite of the distant phylogenetic relationships between the two species. Taken together, these data provide strong evidence to support the hypothesis that species having 31 chromosomes per haploid genome basically retain the ancestral karyotype of Lepidoptera.

### Conclusion

In order to establish new markers for a FISH-based chromosome map of *O. nubilalis* by efficient screening of *Ostrinia* orthologues of *B. mori* genes located on targeted chromosomal regions lacking markers, we determined and assembled 511 Mb genome sequence and constructed a fosmid library of *O. furnacalis.* Consequently, we constructed a cytogenetic map of *O. nubilalis* consisting of 123 BAC and eight fosmid probes covering all 31 chromosomes. This map can be applied to other *Ostrinia* moths and related species, and will be useful for the detection of local chromosomal rearrangements which have occurred uniquely in particular species. The targeted selection of BAC and fosmid clones reported here is especially effective for rapid determination of extensive genomic sequences or genome-wide characterization of multiple-copy genes in organisms lacking genome reference sequences.

We estimated the boundaries between ancient chromosomes on *B. mori* chromosomes 11, 23 and 24 by mapping *O. nubilalis* orthologues, which agreed well with a previous estimation based on sequence comparison between *B. mori* and *M. cinxia.* These findings strongly suggest that lepidopteran species having 31 chromosomes per haploid genome basically retain the ancestral karyotype of Lepidoptera.

## Acknowledgements

We are grateful to Dr. M. R. Goldsmith for critical reading of the manuscript. We thank Aya Kawai and Noriko Fujikawa-Kojima for their technical assistance in FISH analysis. A part of this study is financially supported by the Program for Promotion of Basic Research Activities for Innovative Biosciences (PROBRAIN) and grants 22380040, 23248008, 23380030, 25292204 of Japan Society for the Promotion of Science (JSPS).

## Supplemental data

**Table S1.** Sequences of PCR primers used in this study.

**Table S2.** *O. nubilalis* BAC clones used as probes for FISH analysis shown in Figure 1.

**Table S3.** Summary of genome sequencing and assembly of *O. furnacalis.*

**Table S4.** Mapping of genes and markers whose *B. mori* orthologues are located near the boundaries between ancient chromosomes in previous studies.

**Table S5.** Genotyping of 24 BF1 progeny between *O. nubilalis* and *O. scapulalis.*

